# 12-Hydroxystearic acid induces epidermal keratinocytes to secrete antimicrobial peptides that are potent inhibitors of viral infection

**DOI:** 10.1101/2025.07.01.662536

**Authors:** Obed Manglalsiem Gangte, Krittika Nandy, Tanay Bhatt, Ananya Grewall, Akshata Kulkarni, Vaishali Garg, Anupama Ashok, Sayandip Mukherjee, Morris Wasker, Sandip Pathak, Nitish Kumar, Vibhav Sanzgiri, Harshinie Jayasekera, Naresh Ghatlia, Amitabha Majumdar, Colin Jamora

## Abstract

Epidermal keratinocytes produce antimicrobial peptides (AMPs) that serve as a crucial component of the skin’s innate immune barrier. These peptides effectively target a broad spectrum of pathogenic microorganisms while preserving commensal microbiota essential for skin barrier homeostasis and overall skin health. Regulating the release of these AMPs presents a promising approach to enhancing the skin barrier’s defense mechanisms with minimal side effects. We have identified 12-hydroxystearic acid (12-HSA) as a potent stimulator of AMP secretion from primary epidermal keratinocytes. Mechanistic investigations revealed that, akin to bacterial stimulation, 12-HSA induces AMP release through the downregulation of caspase-8, which subsequently activates the inflammasome. Notably, we discovered that 12-HSA mediates caspase-8 downregulation via the acute activation of DNA methyltransferase 3A (DNMT3A), leading to transcriptional silencing of the caspase-8 locus. Importantly, 12-HSA is widely utilized in the cosmetic industry for its several skin beneficial properties including hydration and emolliency. Our findings suggest that this compound can be leveraged to enhance innate immune defenses in the skin, effectively mobilizing stored AMPs from keratinocytes to counteract microbial threats. This discovery highlights the potential for 12-HSA as a novel agent in dermatological applications aimed at fortifying skin barrier immunity.

## INTRODUCTION

The skin is the main physical barrier between the body and the external environment. In addition to this physical barrier, skin is also an immunological barrier with both innate and adaptive immune systems poised to rapidly mobilize in the event of a barrier breach. Not only is there a robust representation of a large repertoire of myeloid cells and lymphocytes, but the epidermal keratinocytes themselves harbor important immune functioning molecules. In addition to being a rich source of cytokines (Lee et al., 2009), they are a large reservoir of antimicrobial peptides (AMPs), which are a class of small peptides that widely exist in nature; furthermore, these AMPs are an important part of the innate immune system of different organisms. Interestingly, the AMPs produced in the epidermis are stored in lamellar bodies of the differentiated keratinocytes found in the uppermost layer of live cells in the epidermis (the granular layer). Consequently, their release from cells at the interface of the tissue and the environment renders them the first line of defense against pathogenic invasion. AMPs exhibit a broad spectrum of inhibitory activity against bacteria, fungi, parasites, and viruses. Beyond these antimicrobial effects, AMPs also play diverse roles in immune regulation: they can modulate pro-inflammatory responses, recruit immune cells, stimulate cell proliferation, enhance wound healing, alter gene expression, and even target cancer cells. Through these functions, AMPs contribute to immune defense in human skin, respiratory infections, and inflammatory diseases. (de la Fuente-Núñez et al., 2017). Despite their significant biomedical promise, the clinical use of synthetic AMPs has been limited by technical obstacles, including the need to reduce cytotoxicity, improve stability against proteases, and develop cost-effective production methods. Since epithelial barrier tissues like the skin contain a rich reservoir of biologically active AMPs, strategies to selectively upregulate and release these naturally produced peptides on demand could offer a substantial breakthrough in addressing these challenges.

12-Hydroxystearic acid (12-HSA), a hydroxylated fatty acid that is readily derived from its precursor - ricinoleic acid, which is abundantly found in natural fats and oils—most prominently castor oil—is gaining increasing attention in this context. Long valued in the cosmetic and personal care industries for its emolliency, hydration, melanin management and other skin benefits as well as formulation stabilizing properties, 12-HSA is now being re-evaluated for its potential therapeutic effects. Emerging evidence suggests that 12-HSA may bolster skin barrier function and modulate cutaneous immunity (Yarova et al., 2022). However, the molecular and cellular mechanisms through which 12-HSA exerts these effects remain incompletely understood.

Given the rising prevalence of antibiotic-resistant bacteria and the persistent threat of viral infections, enhancing the skin’s innate defenses has become a strategic priority. Strengthening both the physical structure of the epidermis and the underlying immune network could provide a powerful, non-invasive approach to mitigating pathogen entry and systemic infection. In this context, elucidating the functional impact of 12-HSA on skin health may reveal promising avenues for both preventative and therapeutic interventions.

## RESULTS

### 12-HSA induces the expression and secretion of AMPs from keratinocytes

In order to evaluate the potential impact of 12-HSA on skin in an unbiased fashion, we assayed for the effect of this fatty acid on the transcriptional profile of epidermal keratinocytes. Since the differentiated cells of the granular layer play a major role in the barrier function of the skin, we modeled this state of the epidermal keratinocytes in vitro. Moreover, this granular layer is the population of epidermal keratinocytes that would receive the highest exposure of 12-HSA following topical application to the skin. An immortalized keratinocyte cell line (Ramirez et al., 2003) was cultured in low Ca^2+^ medium as previously described (Nowak & Fuchs, 2009). Upon reaching 100% confluency, differentiation was induced by replacing the medium with high Ca^2+^ (1.8 mM) medium for at least 48 hours. Cells were treated with vehicle control or 40 μM 12-HSA followed by RNA extraction and RNAseq analysis to generate a transcriptional profile of the cells. Interestingly, several antimicrobial peptides and related genes were upregulated (Figure 1A). This prompted us to perform a gene of interest (GOI) search where we found that the AMP genes KNG1, RNASE7, CXCL13, HAMP and CALCB were significantly upregulated (Figure 1B). Given the representation of AMPs in this dataset, we assessed whether other well-characterized AMPs produced by differentiated epidermal keratinocytes are upregulated as well in response to 12-HSA. Using qPCR, we observed that the genes of two members of the beta defensin class of AMPs were also upregulated upon treatment with 12-HSA – hBD1 and hBD4 (Figure 1C). Human beta defensins are arguably the most important AMPs in epithelial tissues (Prado-Montes de Oca, 2010). They are known for their broad-spectrum antimicrobial activity due to their non-specificity of targets (Chen et al., 2007; Hancock & Diamond, 2000; Park et al., 2018; Prado-Montes de Oca, 2010; Smiley et al., 2007; van Kilsdonk et al., 2017; Wilson et al., 2013).

**Figure 1.**
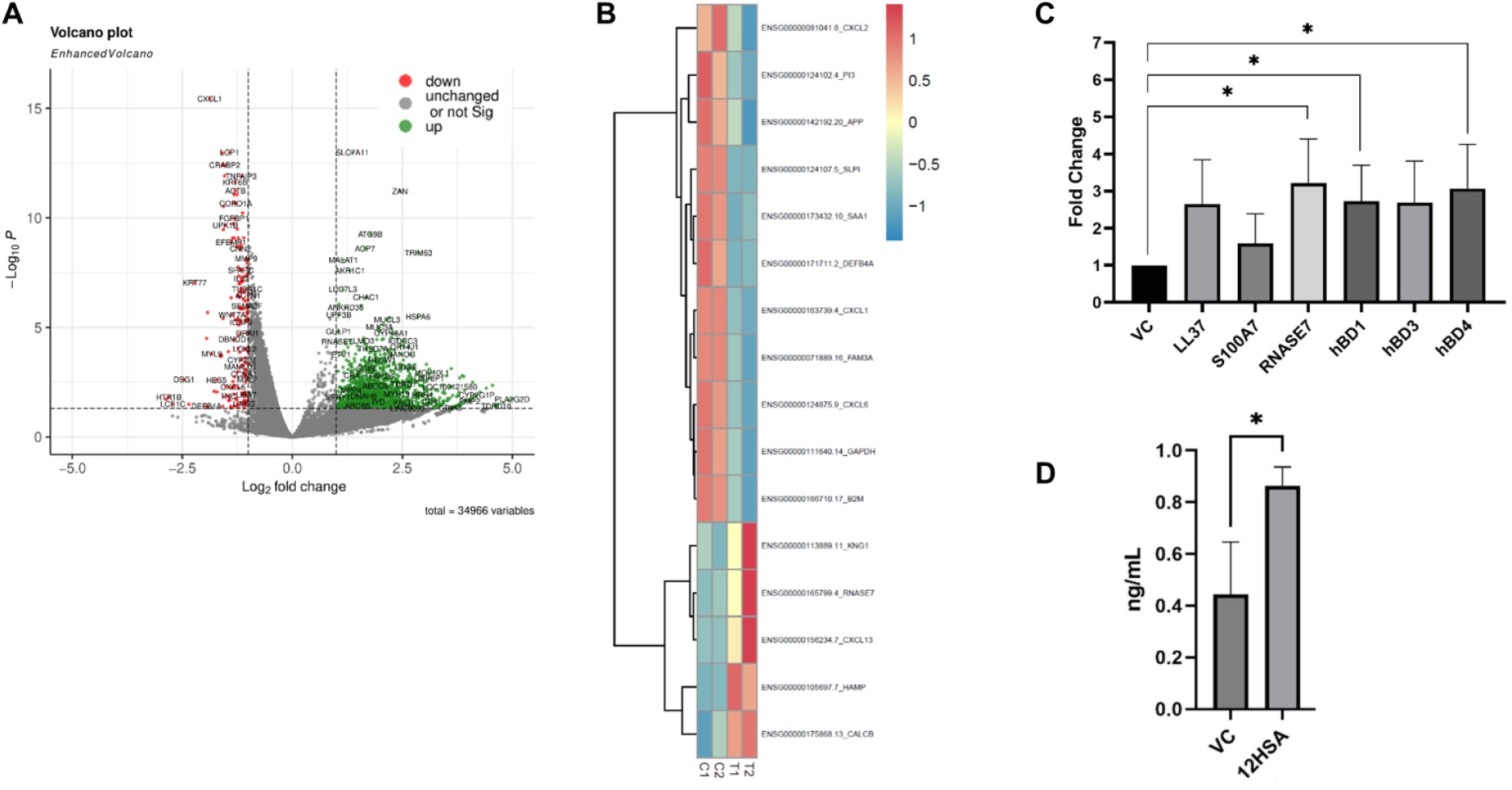
Treatment of kératinocytes with 12-HSA induces upregulation in the expression of AMPs._ (A) Volcano plot of upregulated and downregulated genes. (B) Heatmap of differentially regulated AMP related genes in control versus 12-HSA treatment. (C) RTqPCR of epidermal AMPs; (D) ELISA of LL-37 from conditioned media of cells treated with vehicle control or 12-HSA. * denotes p<0.05.

Given the fact that large reservoirs of AMPs are stored in the keratinocytes of the granular layer, we next examined whether 12-HSA would also induce the secretion of these peptides (or merely increase the intracellular pool stored in lamellar bodies). Analysis of conditioned media from keratinocytes treated with 12-HSA or vehicle control by ELISA revealed that the human cathelicidin LL-37 was elevated in the secretome of cells treated with the fatty acid (Figure 1D). Altogether, these analyses reveal that 12-HSA is capable of upregulating the expression and secretion of AMPs from differentiated epidermal keratinocytes.

### Stimulation of AMP release by 12-HSA is through the downregulation of caspase-8

Previously we reported that the secretion of AMPs from differentiated epidermal keratinocytes in response to exposure to bacteria/pathogen associated molecular patterns (PAMPs) is achieved through the downregulation of caspase-8 (Bhatt et al., 2019). This is in line with the growing appreciation of unconventional roles of caspase-8 apart from its critical function in the extrinsic apoptotic pathway (Lee et al., 2009). In fact, the downregulation of caspase-8 is sufficient to induce a wound healing phenotype in the skin, wherein AMPs are also known to be secreted to protect against microbial invasion through the breached barrier caused by damage to the epidermis (Lee et al., 2009). Moreover, RNAi-mediated reduction in caspase-8 levels in epidermal keratinocytes is sufficient to release the antimicrobial peptide S100A7 (psoriasin) from intracellular stores (Bhatt et al., 2019). We thus examined whether the 12-HSA induced secretion of AMPs in keratinocytes likewise utilizes the downregulation of caspase-8. Analysis of RNA levels by qPCR of cells treated with 12-HSA for 24 hours exhibited a ∼50% decrease in caspase-8 RNA (Figure 2A). We then assessed whether this downregulation was reflected at the protein level. Interestingly, though the RNA is reduced within 24 hours of 12-HSA treatment, the protein level of caspase-8 is only significantly reduced after 48 hours (Figure 2B). Our previous studies demonstrated that the downregulation of Caspase-8, triggered by a wound, activates the inflammasome complex (Lee et al., 2009). Interestingly, this same inflammasome complex was subsequently found to facilitate the secretion of the antimicrobial peptide S100A7 in response to a pathogen associated molecular pattern stimulus (Bhatt et al., 2019). Based on these findings, we investigated whether 12-HSA-induced AMP secretion also engages these inflammasome-related proteins—specifically, p38 MAPK (Figure 2C).

**Figure 2.**
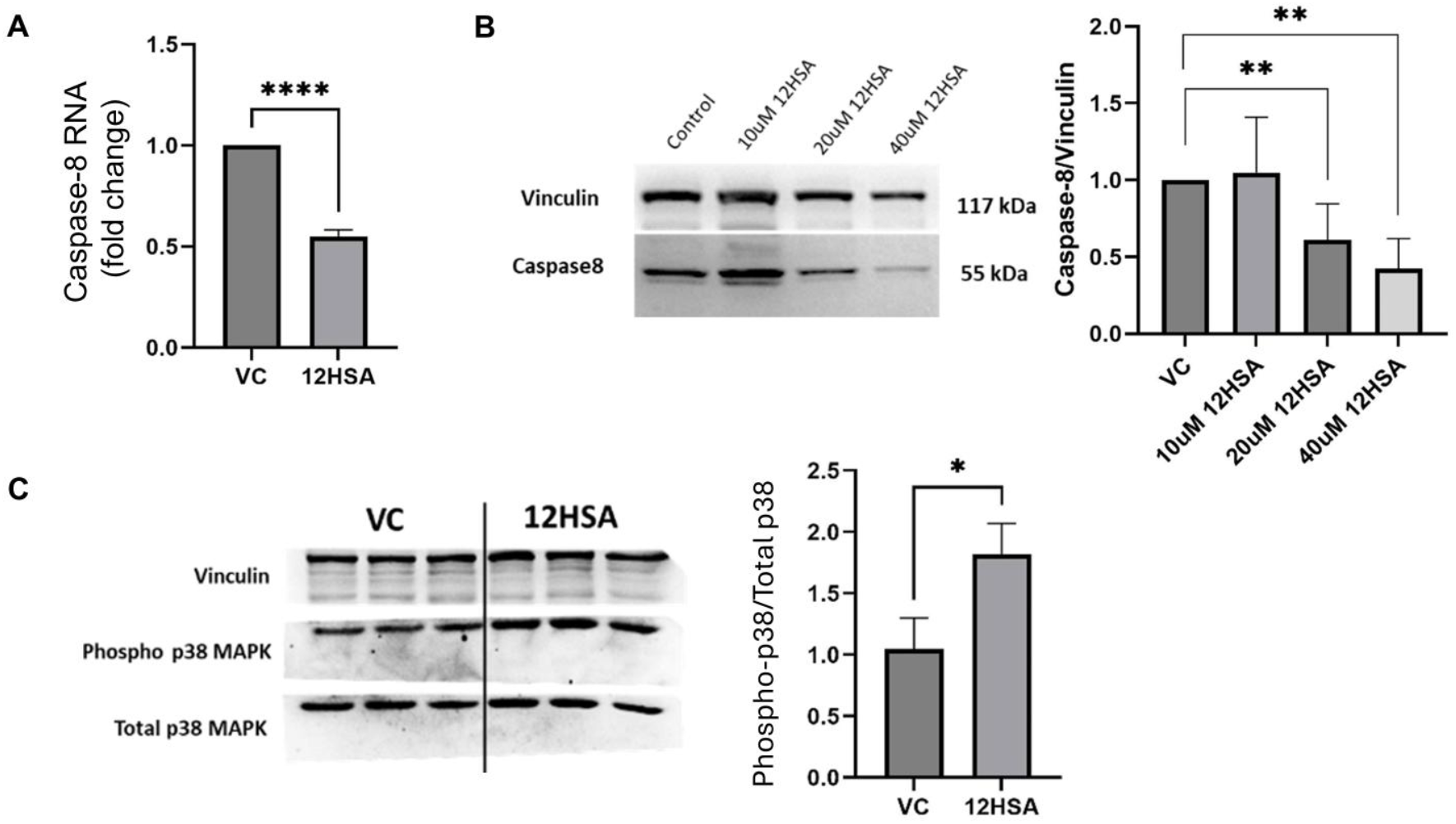
12-HSA induces the downregulation of Caspase-8 which in turn leads to the secretion of AMPs. (A) Fold change of Caspase-8 mRNA levels post 24h of 12-HSA treatment analysed using qPCR technique. (B) Western blot of Caspase-8 protein levels 48h post treatment with 12-HSA. (C) Phosphorylation slate of p38 MAPK 48h post treatment with 12-HSA, analysed using Western blot.

### 12-HSA mediated downregulation of caspase-8 is through transcriptional silencing by DNMT3a

The expression profile of caspase-8 described above suggests that 12-HSA reduces caspase-8 levels by transcriptional silencing within a timeframe of 24 hours, but it that takes longer for this decrease to be manifested at the protein level owing to its relatively long half-life (Bhatt et al., 2019). Drawing from the mechanism of caspase-8 transcriptional repression in the wound healing scenario, we determined whether epigenetic regulation of this locus likewise plays a role in epidermal keratinocytes treated with 12-HSA. Upon scratch wounding of differentiated keratinocytes in vitro, the cells at the leading edge of the wound express the de novo DNA methyltransferase, DNMT3a (Bhatt et al., 2022). In differentiated epidermal keratinocytes, DNMT3A is found primarily in the cytoplasm (Figure 3A). Upon treatment with 12-HSA for 6 hours, there is a substantial amount of DNMT3a localized in the nucleus. The effect of 12-HAS in inducing the nuclear localization of DNMT3a was transient as the protein was found predominantly in the cytoplasm after 20 hours of treatment (Figure 3A). We then investigated whether the nuclear DNMT3a was required for the downregulation of caspase-8. Addition of the 5-azacytidine, a pharmacological inhibitor of methyltransferase activity, with 12-HSA effectively abrogated the reduction of caspase-8 RNA (Figure 3B). In summary, the 12-HSA induced downregulation of caspase-8 RNA is mediated by the nuclear translocation of DNMT3a, which in turn methylates and silences the caspase-8 promoter.

**Figure 3.**
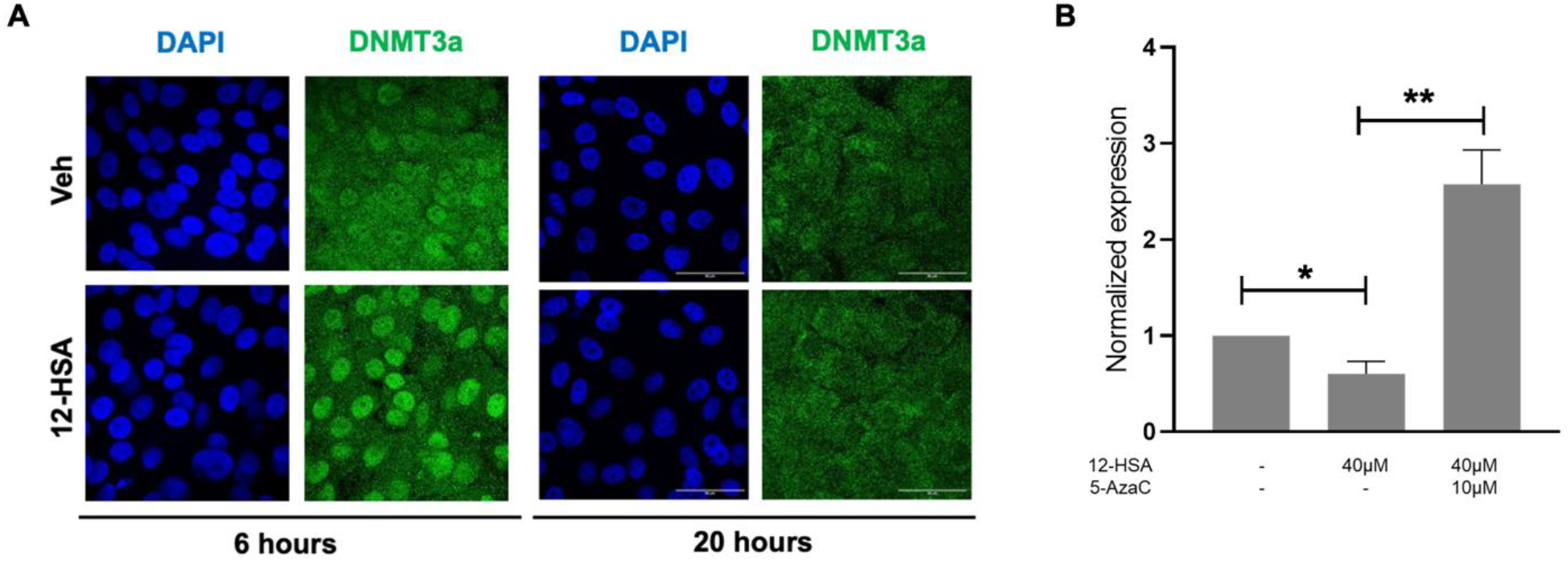
12-HSA mediated downregulatlon of caspase-8 Is through the transcriptional silencing by DNMT3a. (A) lmmunofluorescence of DNMT3a (green) and DAPI (blue) to mark the nucleus of epidermal keratinocytes treated with vehicle control or 12-HSA for 6 hours or 20 hours. (b) Caspase-8 mRNA levels in differentiated keratinocytes treated with 12-HSA and/or DNMT3 inhibitor 5-azacytidine. Error bars represent mean ± SEM. ^*^= p<0.05, ^* *^= p<0.001. Statistical significance was determined by unpaired Student’s *t* test.

### AMPs released from epidermal keratinocytes and skin explants by 12-HSA demonstrates antiviral activity

We reasoned that the ability of 12-HSA to induce AMP release from keratinocytes should endow their secretome with anti-microbial activity. In order to test whether 12-HSA is a viable method of inducing AMPs from a stratified epithelial such as the epidermis, we tested the effect of this fatty acid on human skin explants ex vivo. 10mm punch biopsies were treated with 40μM of 12-HSA or the corresponding volume of vehicle control for 24 hours. Following this treatment, RNA was extracted from the tissue and processed for qPCR analysis of various AMPs. Similar to the effect observed in vitro, 12-HSA treated skin explants exhibited a significant increase in the expression of beta defensin, RNase7 and LL-37 (Figure 4A). Analysis of the conditioned media of these explants by ELISA revealed that there was a corresponding increase in the amount of RNase7 and LL-37 peptides secreted in response to 12-HSA (Figure 4B). In vitro data implicates the downregulation of caspase-8 mRNA as a stimulus for the release of AMPs (Figure 2A and B) (Bhatt et al., 2019). In line with these observations, we have also found that caspase-8 mRNA levels are reduced ∼50% upon treatment of epidermal explants with 12-HSA (Figure 4C).

**Figure 4.**
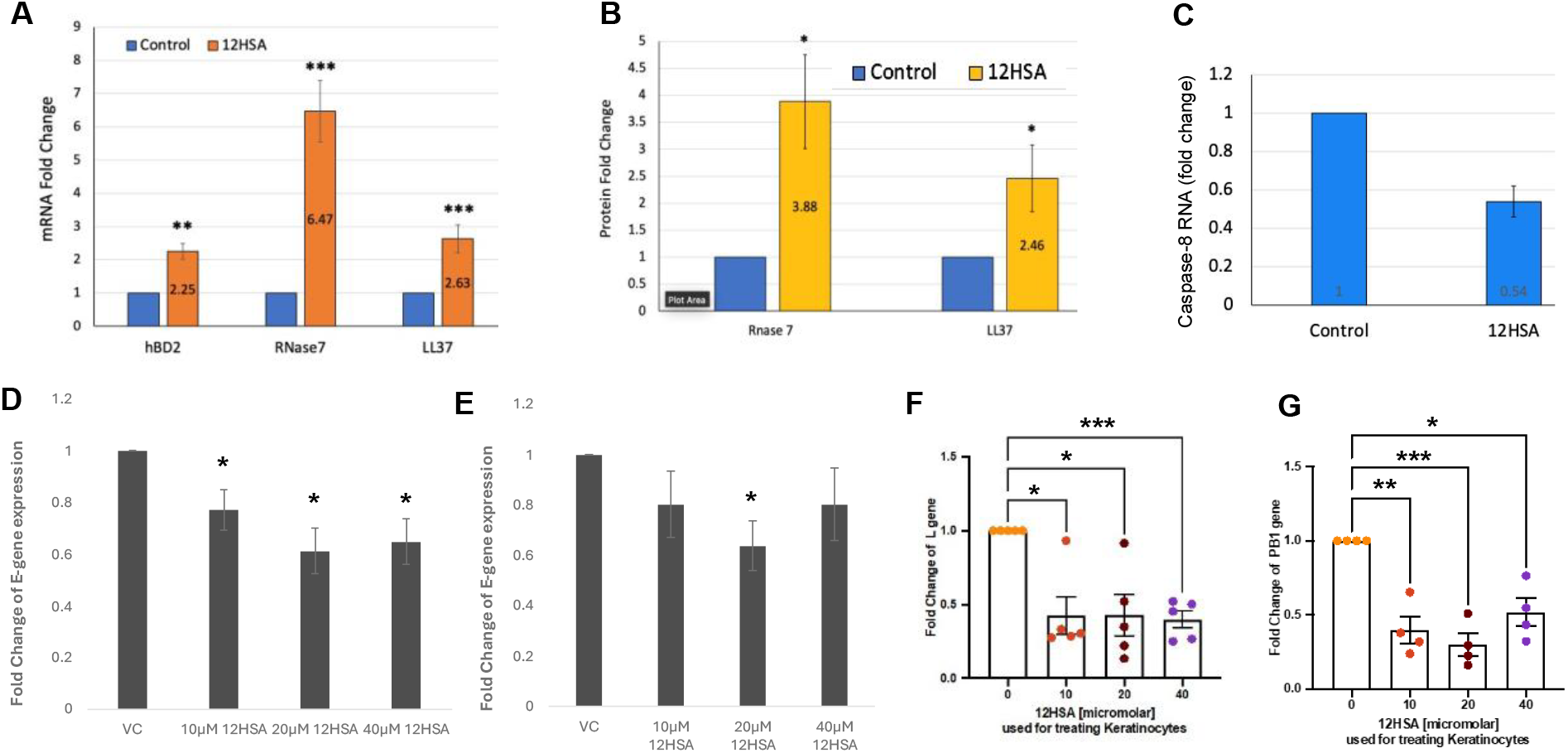
12-HAS mediated secretion of AMPs can inhibit viral infection. (A) PCR analysis of AMP mRNA expression in keratinocytes treated with vehicle control (blue) or 12-HSA (orange) (B). old change of AMP secretion measured by ELISA of conditioned media from keratinocytes treated with vehicle control (blue) or 12-HSA (yellow) (C) Caspase 8 RNA levels in epidermal keratinocytes treated with vehicle control or 12-HSA (D),(E) Amount of the E-gene from the SARS Co V-2 virus detected in Calu3 cells by PCR following treatment of skin explants and primary keratinocytes respectively with 12-HSA (F) Amount of L gene from RS detected in in Calu3 cells by PCR following treatment of skin explants with vehicle control or increasing concentration of 12-HSA. (G) PB1 gene of IVA detected in Calu3 by PCR post treatment of skin explant with 12HSA. ^*^= p <0.05, ^* *^ = p = <0.001, ^* * *^ = p <0.0001.

Previously we reported that a recombinant AMP, LL-37/cathelicidin, was effective against multiple strains of SARS-CoV-2, the virus causing COVID (Bhatt et al., 2023). The broad spectrum of activity against viral variants was owed, in part, to the ability of LL-37 to disrupt the viral membrane, thereby making it agnostic to any protein mutations. Given recent reports of the resurgence of COVID infections in Southeast Asia, new avenues of mitigating infections regardless of subtypes of the virus remains a pressing need. We therefore investigated whether the AMPs released from the skin in response to 12-HSA, are capable of hindering SARS-CoV-2 infection. Interestingly, though keratinocytes express the ACE2 receptor, thereby theoretically rendering them susceptible to infection, skin contact is not the normal route of transmission. We therefore applied 12-HSA (or vehicle control) to the surface of the skin explants and incubated for 24 hours. Following this, the secretome from the epidermis of the skin was collected and incubated with SARS-CoV-2 and following 1 hour of treatment the virus was added to the infection competent cell line Calu3. Infection was measured by qPCR analysis for the viral E-gene amongst the RNA extracted from the Calu3 cells. Interestingly, the secretome released by 12-HSA treated skin, likely mediated through AMP related pathways, exhibited a ∼40% decrease in infection relative to the vehicle control treated virus (Figure 4D). Similarly, differentiated primary human keratinocytes were also treated with 12-HSA for 24 hours after which the secretome was incubated with SARS-CoV-2 as described previously. The secretome from 12-HSA treated primary keratinocytes also exhibited a ∼40% decrease in infection relative to the vehicle control treated virus (Figure 4E).

Respiratory Syncytial Virus (RSV) is a primary cause of acute lower respiratory tract infections, particularly affecting young children (Gatt et al., 2023) and posing risks such as severe pneumonia and bronchitis in adults with chronic conditions and individuals over 65(Savic et al., 2023). The recent SARS-CoV-2 pandemic has significantly disrupted RSV’s seasonal patterns, potentially due to immune degradation caused by SARS-CoV-2, increasing susceptibility in patients (Abu-Raya et al., 2023). With rising concerns over comorbidity and mortality, exploring 12-HSA-induced AMPs as a strategy against multiple respiratory viruses could potentially be an interesting approach. As such, we conduct a similar experiment to the one described above and found that RSV showed a ∼60% reduction in relative infection (Figure 4F), indicating a potential broad antiviral activity of 12-HSA-induced AMPs.

Viral co-infections are also highly common during the influenza season leading to severe clinical illness (Xiang et al., 2021). Influenza A virus (IAV) had been shown to co-infect a number of SARS-CoV-2 infected patients during the COVID-19 pandemic and significantly degrading patients’ conditions and increasing the risk of death (Goka et al., 2013). We therefore reasoned that 12-HSA mediated intervention could be a promising approach. Secretome from 12-HSA treated skin explants were used to treat IAV, similar to the process described above. The PB1 gene of IVA showed a significant reduction of ∼70% relative to the vehicle control treated virus (Figure 4G). Altogether, these results indicate that 12-HSA can be an effective means of inducing AMP secretion from the skin that are bioactive against respiratory viruses.

## DISCUSSION

We have found that a common component in cosmetic and personal care products, 12-hydroxystearic acid (12-HSA) can also be utilized to enhance the skin defense barrier by enhancing the expression and secretion of antimicrobial peptides stored in epidermal keratinocytes. Interestingly, the mechanism of action of 12-HSA appears to converge on signaling pathways that stimulate cutaneous AMP release in the context of wound healing and exposure to PAMPs. Among the components upstream in this signaling hierarchy is the protein caspase-8, whose downregulation is sufficient to induce AMP secretion. Similar to the wound-healing scenario, the reduction of caspase-8 expression is accomplished through the acute and short-termed transcriptional silencing by the epigenetic regulator DNMT3a. Release of cellular tension upon wounding has been demonstrated to induce nuclear translocation of DNMT3a, but whether a fatty acid such as 12-HSA is capable of releasing the skin tension in the cell remains to be determined.

The topical application of 12HSA to the skin surface to enhance skin barrier health via AMP release would be a useful modality in enhancing the immunological, infection protection. A next question would be to determine whether other epithelial barriers such as the lungs and gut can also respond to 12-HSA exposure to prevent respiratory and gastrointestinal infections, respectively.

## ACKNOWLEDGEMENTS

The authors would like to thank the members of Jamora laboratory for their critical review of the work and insightful discussions. Funding for this project was provided by Unilever Pvt. Ltd. and core funds from inStem, Bangalore. This research work was carried out in part at the DST-FIST Confocal Microscope Facility (R-250), a part of the Core Imaging Facility (CIF) of the Department of Life Sciences, SNS, which is managed by the Shiv Nadar IoE DTU Delhi-NCR and is funded by the Department of Science and Technology (Grant No. SR/FST/LS-1/2017/59(c)

## MATERIALS AND METHODS

### Approvals

All experimental work was approved by the inStem Institutional Biosafety Committee and the Shiv Nadar Institution of Eminence Biosafety Committee. Informed consent was obtained from all patients for skin sample collection and experimentation and study protocols were approved by the Institutional Human Ethics Committee of inStem.

### Cell culture and Treatments

The human keratinocyte cell line Ker-CT (ATCC CRL-4048) was cultured in low calcium media comprising of EpiLife (GIBCO #MEPI500CA) medium with growth supplement (HKGS, GIBCO #S0015). The media was further supplemented with 1.2% calcium chelated serum and, 100U/ml penicillin and 100μg/ml streptomycin (Gibco) and grown at 37°C in a 5% CO2 incubator. The cells were allowed to grow to 100% confluency and then differentiated by increasing the calcium concentration to 1.8 mM in EpiLife without any other supplementation for 48 hours. The differentiated cells were then treated with different concentrations of 12-HSA (10μM, 20μM and 40μM) which was provided by Unilever, for 24 or 48 hours depending upon the experiment.

### Transcriptomics

RNA was extracted from Ker-CT cells treated with either Control (ethanol) or 20μM 12-HSA for 24 hours (two biological replicates per condition). The quality of RNA samples was evaluated using the Agilent TapeStation with High Sensitivity RNA ScreenTape, and samples with a RIN score exceeding 5 were chosen for library preparation. mRNA was enriched using oligo-dT beads via the Dynabeads mRNA Purification Kit (Invitrogen). The enriched mRNA was subsequently utilized for library preparation with the MGIEasy RNA Library Prep Kit (MGI, Cat No: 1000005276). Libraries underwent PCR amplification and circularization using the MGIEasy Circularization Module (MGI, Cat No: 1000005260). These circularized libraries were converted into DNA Nanoballs (DNBs) and sequenced on the DNBSEQ-G400RS platform utilizing FCL PE100 flow cell chemistry (1000016949 and 1000016985). Each sample yielded approximately 30–50 million reads. Analysis of the data was outsourced to a commercial facility.

### Quantitative Real-Time PCR

RNA was isolated from the human keratinocyte cell line Ker-CT using RNAiso Plus (Takara). PrimeScript kit (Takara) was used to prepare cDNA using 1 μg of isolated RNA. Quantitative PCR (qPCR) was then done with cDNA equivalent to approximately 100 ng of RNA using the SsoFast 2x master mix (BioRad). All reactions were performed in technical triplicates using the CFX384 Touch Real-time PCR detection system (BioRad). TATA-binding protein (TBP) expression was used as reference and normalization for all reactions. The sequences of primers used are given below:

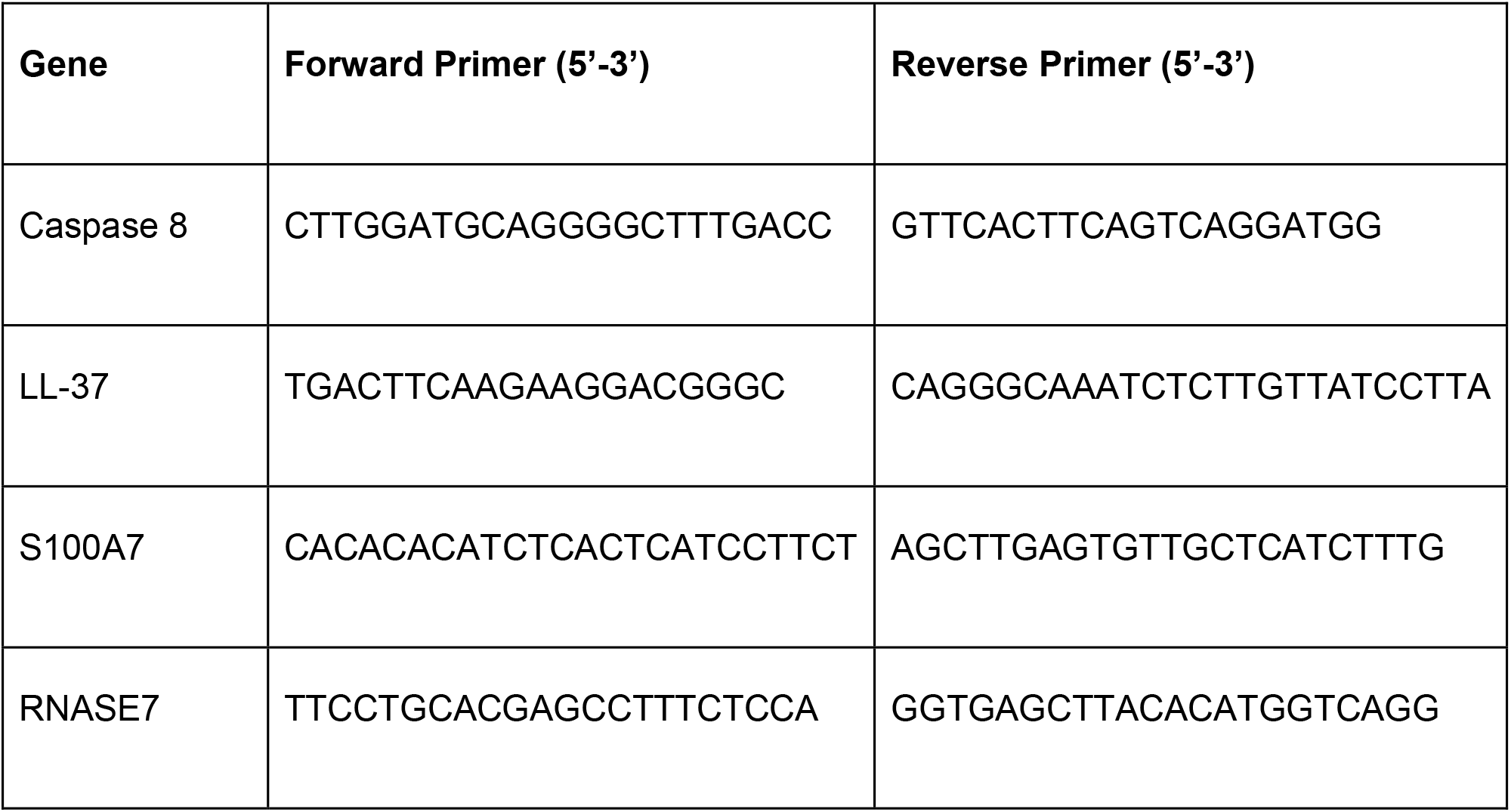
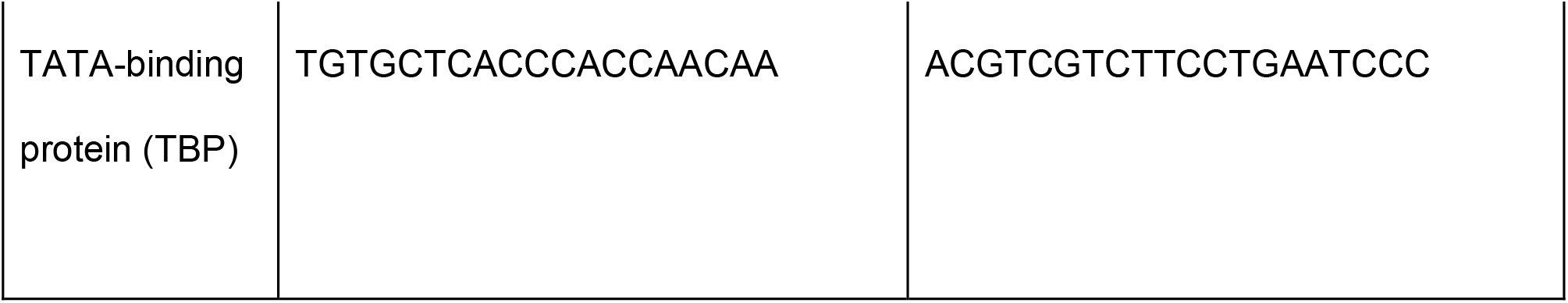

### Human Skin Explant Treatments

The skin tissues were transported in Dulbecco’s Modified Eagle Medium (DMEM) supplemented with 10% fetal bovine serum (FBS) and 2x Anti-Anti (Thermo). Upon arrival, the tissues were washed twice with 1x PBS for 30 min each. Excess fatty tissues were removed, and the skin was placed and spread in sterile 100 mm culture dishes with dermis on the lower side. The epidermal side is gently wiped with autoclaved lint free tissue paper. The tissues were cut into equal sized pieces (10×10 mm) and transferred to the 12-well plate, the dermis was allowed to stick to the bottom of the plate for 10 min at 25’ C. 500 uL of basal Epilife media was added around the tissue in a way that the cut sides of the skin tissue touch the media and epidermis remain in contact with air.

Specific concentration of 12HSA was prepared in PBS and 30ul drop was gently spread over the epidermis. After 24 hr the epidermis was washed with 1x PBS and 12HSA treatment was repeated for another 24 hr. After 48 hr of 12HSA treatment, the spent media was collected and tested for various AMPs. Some treated explants will be taken for trizol based RNA isolation and assessment of AMPs gene expression. The remaining explants received equal quantities of SARS-CoV-2 particles (within 10ul volume) spread on the epidermal surface and incubated for 30 min at 37’ C. These skin explants were then washed using 250 ul of DMEM to collect the virus particles. These washed particles were incubated with Calu3 cells for 1 hr for adsorption. After 24 hr, the Calu3 cells were lysed in trizol and SARS-CoV-2 were quantified through E-gene based qPCR and normalised using TBP.

### Western blot analysis

The lysates of the human keratinocyte cell line Ker-CT were prepared in RIPA buffer with protease inhibitor (Sigma, #P2714) and PhosSTOPTM (Sigma, # 12352204) where required. The lysate was the sonicated at 4°C. The protein lysates were then mixed with 4X Laemmli buffer and heated at 95°C for 5 minutes before loading then in 10% polyacrylamide gel. The proteins were then transferred on to a nitrocellulose membrane which was then blocked using 5% Blotto (Santa Cruz Biotechnology, sc-2325) in Tris buffer Saline containing 0.1% Tween (TBST) or 5% BSA for phosphorylated proteins for 60 minutes. The blots were probed overnight with the respective primary antibodies viz. Caspase-8 (Cell Signalling, #9746S), p38 MAPK (Cell Signalling, #9212S), Phospho p38 MAPK (Cell Signalling, #4511S) and Vinculin (Sigma, #V9264). After washing with TBST, they were then probed with HRP tagged secondary antibodies and washed again. Enhanced Chemiluminescence substrate (ECL, Merck) was then used to detect the signals using an iBright FL (Thermo) detector. Quantification of the bands was done using Fiji (ImageJ) software and normalized to Vinculin which was used as the loading control.

### ELISA

Conditioned media from differentiated Ker-CT cells were collected 72 hours post treatment with 12-HSA. The concentration of different AMPs was determined by ELISA using commercially available analysis kits - LL-37 (Hycult Biotech, HK321), S100A7 (CircuLex, 10j17A) and RNASE7 (Hycult Biotech, HK371) following the manufacturers’ protocols. A microplate reader (Varioskan, Thermo) was then used to determine the absorbances at 450nm.

### Immunostaining

The differentiated Ker-CT cells post treatment with 12-HSA were fixed in 4% paraformaldehyde (PFA) before permeabilizing them with 0.2% Triton X 100. The cells were then stained overnight with anti-DNMT3a antibody (Santa Cruz, sc-365769) overnight. Alexa Fluor 568–labelled secondary antibodies (Jackson ImmunoResearch) were used at a dilution of 1:400. After which, DAPI or Hoechst stain was used to mark the nucleus. The Nikon A1R Confocal Microscope was then used for imaging and Fiji (ImageJ) software was used for processing the images.

### Keratinocyte Culture, 12HSA Treatment, and Conditioned Media Collection

Commercially available human keratinocytes were cultured to 100% confluence and subsequently differentiated for 48 hours using high-calcium media. Differentiated keratinocytes were then treated with varying concentrations of 12-hydroxystearic acid (12HSA), ranging from 0 µM (vehicle control) to 80 µM. Following a 48-hour treatment period, the conditioned media was collected, flash-frozen, and stored at -80°C until use.

### Antiviral Neutralization Assay

MDCK (for Influenza A Virus, IAV) or VeroE6 (for Respiratory Syncytial Virus, RSV) host cells were cultured in 12-well plates to 75% confluence. For virus neutralization, 80 µL of keratinocyte conditioned media was incubated with 20 µL of either RSV or IAV stock virus at 37°C for 1 hour. This 100 µL virus-conditioned media mixture was then combined with 800 µL of fresh cell culture media and added to the respective host cells after removal of existing media. Following a 1-hour adsorption period at 37°C, the inoculum was removed, cells were washed once with 1X PBS, and fresh cell culture media was added. Infected host cells were then incubated for 18 hours at 37°C with 5% CO2.

### Viral Gene Expression Analysis by RT-qPCR

After the 18-hour incubation, host cells were harvested using NeoDx kit lysis buffer according to the manufacturer’s protocol. Total RNA was isolated, and cDNA was synthesized. Viral gene expression was quantified by qPCR using virus-specific primers (L gene for RSV, PB1 gene for IAV). The cycle threshold (Ct) values were determined for various treatments and normalized to GAPDH expression as an endogenous control.

## Notes

### Competing Interest Statement

The authors have declared no competing interest.

